# Pharmacological complementation remedies an inborn error of lipid metabolism

**DOI:** 10.1101/848119

**Authors:** Meredith D. Hartley, Mitra D. Shokat, Margaret J. DeBell, Tania Banerji, Lisa L. Kirkemo, Thomas S. Scanlan

## Abstract

X-linked adrenoleukodystrophy (X-ALD) is a rare, genetic disease in which increased very long chain fatty acids (VLCFAs) in the central nervous system (CNS) cause demyelination and axonal degeneration, leading to severe neurological deficits. Sobetirome, a potent thyroid hormone agonist, has been shown to lower VLCFA levels in the periphery and CNS. In this study, two pharmacological strategies for enhancing the effects of thyromimetics were tested in *Abcd1* KO mice, a murine model that has the same inborn error in metabolism as X-ALD patients. First, a sobetirome prodrug (Sob-AM2) with increased CNS penetration lowered CNS VLCFAs more potently than sobetirome, and was better tolerated with lower peripheral exposure, but was unable to unable to break the efficacy threshold of CNS VLCFA lowering in *Abcd1* KO mice. Second, co-administration of thyroid hormone with sobetirome enhanced VLCFA lowering in the periphery compared to sobetirome alone but did not produce greater lowering in the CNS. These data suggest that the extent of CNS VLCFA lowering in *Abcd1* KO mice is limited by a mechanistic threshold related to slow turnover kinetics, potentially related to the lack of frank X-ALD disease in this model. However, Sob-AM2 has improved potency at correcting the lipid abnormality associated with X-ALD in the CNS with better tolerance than the parent drug sobetirome.

## Introduction

X-linked adrenoleukodystrophy (X-ALD) is an inborn error of metabolism in which very long chain fatty acids (VLCFAs) are elevated in plasma and tissues resulting in adrenal gland dysfunction and central nervous system (CNS) demyelination. VLCFAs are fatty acids with 22 or more carbons. VLCFA accumulation results from mutations in *ABCD1*, which encodes a peroxisomal transporter that is critical for VLCFA degradation. A proposed therapeutic strategy for X-ALD invokes a second transporter, ABCD2, which has overlapping substrate specificity with ABCD1 (Netik et al., 1999). Genetic upregulation of *ABCD2* results in the reduction of elevated VLCFA levels in *Abcd1* knockout (KO) mice, a mouse model that replicates the elevated VLCFAs in X-ALD (Pujol et al., 2004).

Recent work has demonstrated the success of a pharmacological complementation strategy based on upregulation of *ABCD2* by a thyroid hormone agonist, or thyromimetic (Hartley et al., 2017). Sobetirome, a potent thyromimetic, lowered VLCFAs in *Abcd1* KO mice in disease relevant tissues including adrenal glands, testes, and brain. In peripheral tissues VLCFAs were lowered by ~50%; however, in the brain, VLCFAs were only lowered by ~15-20%. Turnover of many lipids in the brain is slow to the point of being challenging to measure.

Studies in mice have estimated that myelin lipid half-lives range from 30-360 days depending on the age of mice and the lipid identity (Ando et al., 2003). We previously hypothesized that the limited sobetirome-induced VLCFA lowering observed in the CNS was due to the slow kinetics of lipid turnover in the CNS (Hartley et al., 2017). However, it was also possible that pharmacologically-induced VLCFA lowering was limited by how much sobetirome reached the CNS.

This study was designed to address this question by using two different strategies to increase thyroid hormone action in the CNS. The first strategy employed Sob-AM2, an amide prodrug derivative of sobetirome that improves distribution of sobetirome to the CNS by 10-fold and increases the brain-to-serum ratio by 60-fold (Ferrara et al., 2017; Meinig et al., 2017, 2019; Placzek et al., 2016). Sob-AM2 is converted to sobetirome via hydrolysis by fatty acid amide hydrolase (FAAH), which is highly expressed in the CNS (Meinig et al., 2017). Comparing *Abcd1* KO mice dosed with Sob-AM2 to those dosed with sobetirome probes whether increased CNS exposure of sobetirome would lead to increased VLCFA lowering in the CNS.

The second strategy involved correcting the depletion of endogenous thyroid hormones that occurs after chronic sobetirome treatment at the doses required to affect VLCFA lowering in *Abcd1* KO mice. We recently demonstrated that thyromimetics like sobetirome and Sob-AM2 interact with the hypothalamic-pituitary-thyroid (HPT) endocrine axis to suppress thyroid stimulating hormone (TSH), resulting in suppressed levels of thyroxine (T4) and 3,5,3’-triiodothyronine (T3) (Ferrara et al., 2018). This prompted the question of whether depleted levels of endogenous thyroid hormone led to a hypothyroid CNS that countered the thyromimetic effects of sobetirome. If this is the case, restoring thyroid hormone levels to normal might enhance the observed sobetirome-induced VLCFA lowering in the brain and other tissues.

## Results

### Sob-AM2 lowered peripheral and central VLCFAs after oral administration

Sobetirome reduced serum and tissue VLCFAs in *Abcd1* KO mice after oral administration at 80 and 400 μg/kg/day (Hartley et al., 2017). In an effort to expand upon this finding, we performed dose-response experiments for both sobetirome and Sob-AM2. Drug administration was done according to the previous study design by treating *Abcd1* KO mice with chow containing sobetirome or Sob-AM2 for 12 weeks starting at 3 weeks of age. Sobetirome chow was prepared containing 0.04 and 0.13 mg/kg chow (corresponding to daily nominal doses of 9 and 27 μg/kg/day) to add to the previously studied doses (80 and 400 μg/kg/day). In the previous study, we observed weight loss and sporadic, premature death after ~8-10 weeks of treatment at sobetirome doses of 400 μg/kg and above. However, since the prodrug Sob-AM2 has reduced peripheral exposure of the parent drug sobetirome (Meinig et al., 2017), we hypothesized that it should be better tolerated at higher doses. Sob-AM2 chow was prepared at 0.02, 0.05, 0.14, 0.42, 1.56, and 5.12 mg/kg chow (corresponding to daily nominal doses of 3, 9, 28, 84, 312, and 1024 μg/kg/day). The Sob-AM2 doses at 9, 28, and 84 μg/kg corresponded to equimolar doses of sobetirome at 9, 27, and 80 μg/kg. In contrast to chronic dosing with sobetirome at doses greater than 80 μg/kg/day, no weight loss was observed with the higher doses of Sob-AM2 (312 or 1024 μg/kg/day) over 12 weeks (Figure S1), supporting the conclusion that Sob-AM2 has reduced peripheral exposure and less chronic toxicity than sobetirome.

*Abcd1* KO mice have elevated VLCFAs in blood and all X-ALD disease relevant tissues. After 12 weeks of *ad lib* feeding with the prepared drug chows, we isolated serum, adrenal glands, testes, brain, and spinal cord. Two assays were utilized to measure C26:0, which is the predominant biomarker of X-ALD (Kemp et al., 2016). In the first assay, all fatty acid ester derivatives were hydrolyzed and derivatized into pentafluorobenzyl bromide esters. C22:0 and C26:0 were quantified by gas chromatography-mass spectrometry (GC-MS). C22:0 levels were unaffected by treatment, and were used to normalize the data, which is reported as C26/C22. The second assay measured a specific lipid species, C26:0-lysophophatidylcholine (C26-LPC) by liquid chromatography-tandem MS (LC-MS/MS). C26-LPC is a plasma biomarker of X-ALD used for diagnosis and newborn screening (Sandlers et al., 2012). We measured C26-LPC in serum, brain, and spinal cord of *Abcd1* KO mice using a similar LC-MS/MS method.

In peripheral tissues such as serum, adrenal glands, and testes, C26/C22 and C26-LPC were reduced with both sobetirome and Sob-AM2 (Figure 1, Figure S2, Tables S1-S3). The maximal lowering of serum and adrenal glands (Figure 2A) was significantly attenuated with Sob-AM2 (20-30%) relative to sobetirome (50-60%), which is consistent with the reduced peripheral sobetirome exposure afforded by the prodrug strategy. Testes had more robust maximal lowering with Sob-AM2 (~50%), which may be attributed to relatively high levels of FAAH in the testes (Wei et al., 2006). However, the Sob-AM2 VLCFA lowering in testes was less robust than that seen with sobetirome in testes (~60%).

**Figure 1.**
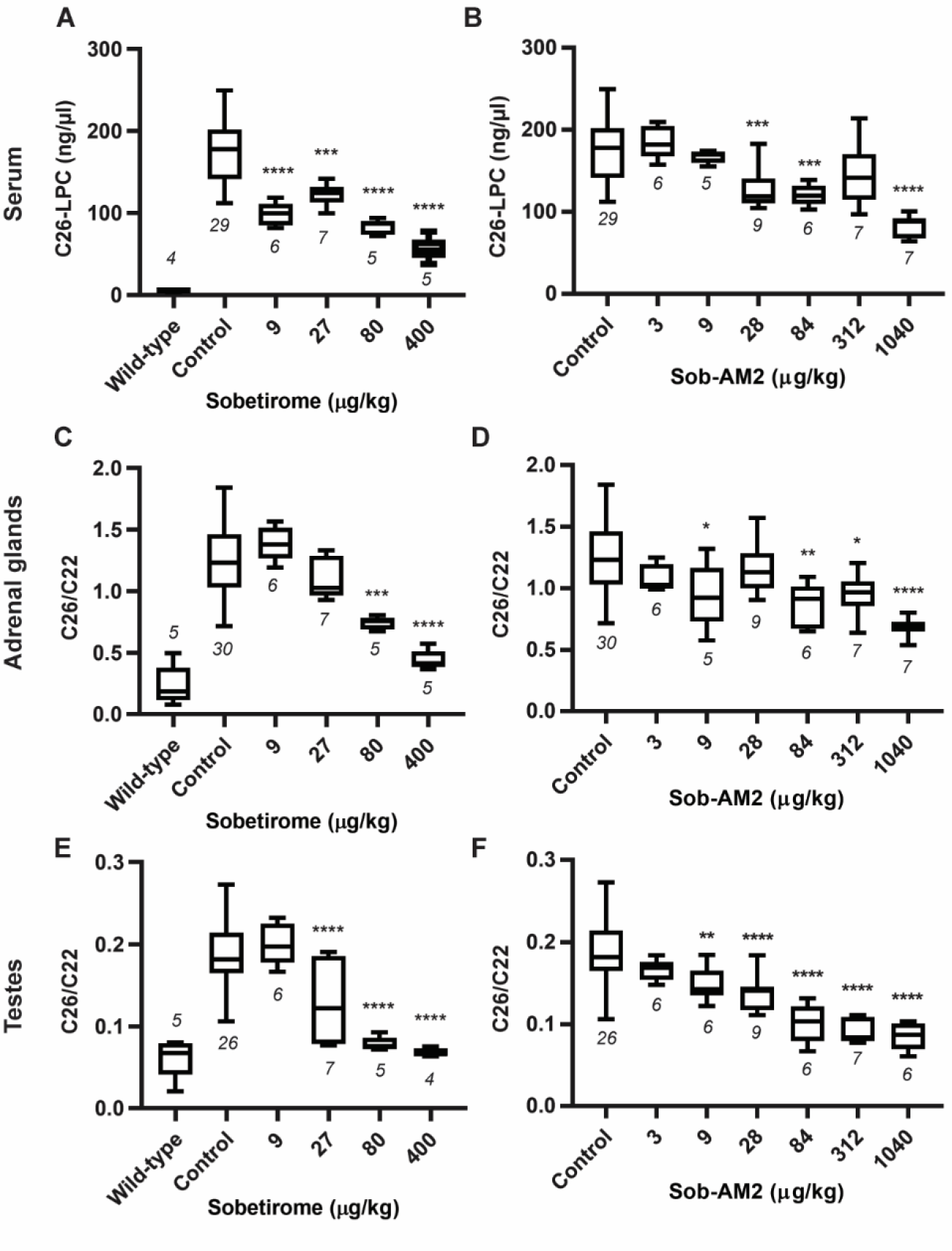
Sobetirome and Sob-AM2 lower C26-LPC and C26/C22 in peripheral tissues. (A – F) Male *Abcd1* KO mice were administered chow containing sobetirome or Sob-AM2 from 3-15 weeks of age. The chow was compounded with sobetirome or Sob-AM2 at the estimated concentration required to administer the daily dose shown in the figure. C26-lysophosphatidylcholine (C26-LPC) was measured by LC-MS/MS in serum (A and B). Total C26 and C22 were measured by GC-MS and the C26/C22 ratio is reported for adrenal glands (C and D) and testes (E and F). All controls for each tissue are combined into a single group. Wild-type mice levels (A, C, and E) were not included in the statistical analyses. All data are represented as box and whisker plots with the error bars representing minimum and maximum. The number of animals is indicated below each dataset in the figure. Statistical analysis was performed using a one-way ANOVA test with Dunnett’s post-test for multiple comparisons between control and each dose (*P < 0.05, **P < 0.01, ***P < 0.001, ****P < 0.0001).

**Figure 2.**
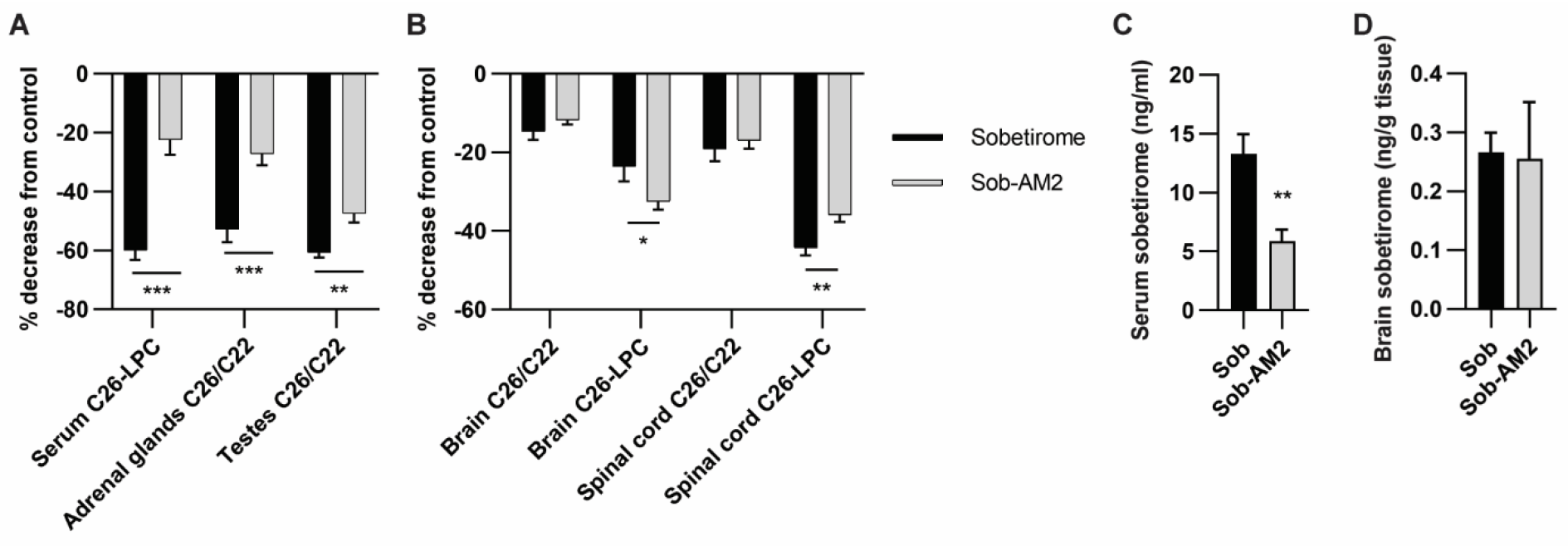
Sob-AM2 lowers VLCFAs in the CNS to the same extent as sobetirome with lower systemic exposure. (A and B) The percent change between the control group and the maximal (or saturating) dose effect is shown for peripheral tissues and CNS tissues. The saturating dose effect was defined as the average of the 80 μg/kg and 400 μg/kg groups for sobetirome, and the average of the 84 μg/kg and 312 μg/kg groups for Sob-AM2. (C and D) Sobetirome levels in serum and brain were measured by LC-MS/MS from mice administered 80 μg/kg sobetirome or 84 μg/kg Sob-AM2. All data are represented as the mean, and the error bars represent SEM. Statistical analysis was performed using an unpaired t-test (*P < 0.05, **P < 0.01, ***P < 0.001).

In both the brain and spinal cord, treatment with sobetirome or Sob-AM2 lowered both C26/C22 and C26-LPC (Figures 3 and 4, Figure S2, Tables S1, S3 and S4). In the brain, the drugs lowered C26/C22 by 12-20% and C26-LPC by 25-30% (Figure 2B). Spinal cord is a critical disease tissue since two-thirds of X-ALD patients develop adult-onset adrenomyeloneuropathy (AMN), which is primarily a disease involving spinal cord pathology (Kemp et al., 2016). The effects of thyromimetics on the spinal cord were not reported previously. Sobetirome and Sob-AM2 maximally lowered C26/C22 by 15-20% and C26-LPC by ~40% (Figure 2B) in the CNS.

**Figure 3.**
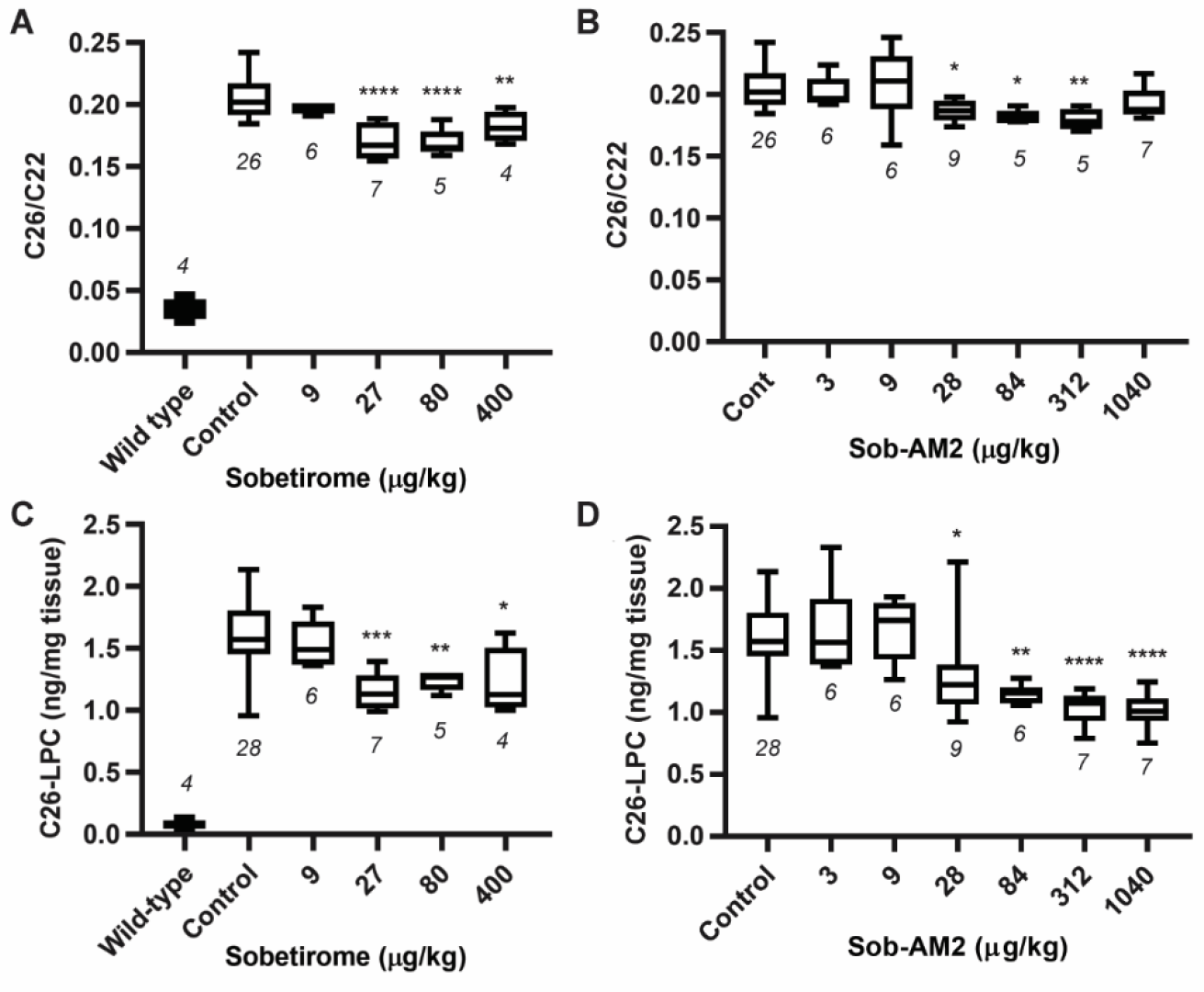
Sobetirome and Sob-AM2 lower C26-LPC and C26/C22 in the brain. (A – D) Male *Abcd1* KO mice were administered chow containing sobetirome or Sob-AM2 from 3-15 weeks of age. The chow was compounded with sobetirome or Sob-AM2 at the estimated concentration required to administer the daily dose shown in the figure. (A and B) Total C26 and C22 in the brain were measured by GC-MS and the C26/C22 ratio is reported. (C and D) C26-lysophosphatidylcholine (C26-LPC) in the brain was measured by LC-MS/MS. All controls for each assay are combined into a single group. Wild-type mice levels (A and C) were not included in the statistical analyses. All data are represented as box and whisker plots with the error bars representing minimum and maximum. The number of animals is indicated below each dataset in the figure. Statistical analysis was performed using a one-way ANOVA test with Dunnett’s post-test for multiple comparisons between control and each dose (*P < 0.05, **P < 0.01, ***P < 0.001, ****P < 0.0001).

**Figure 4.**
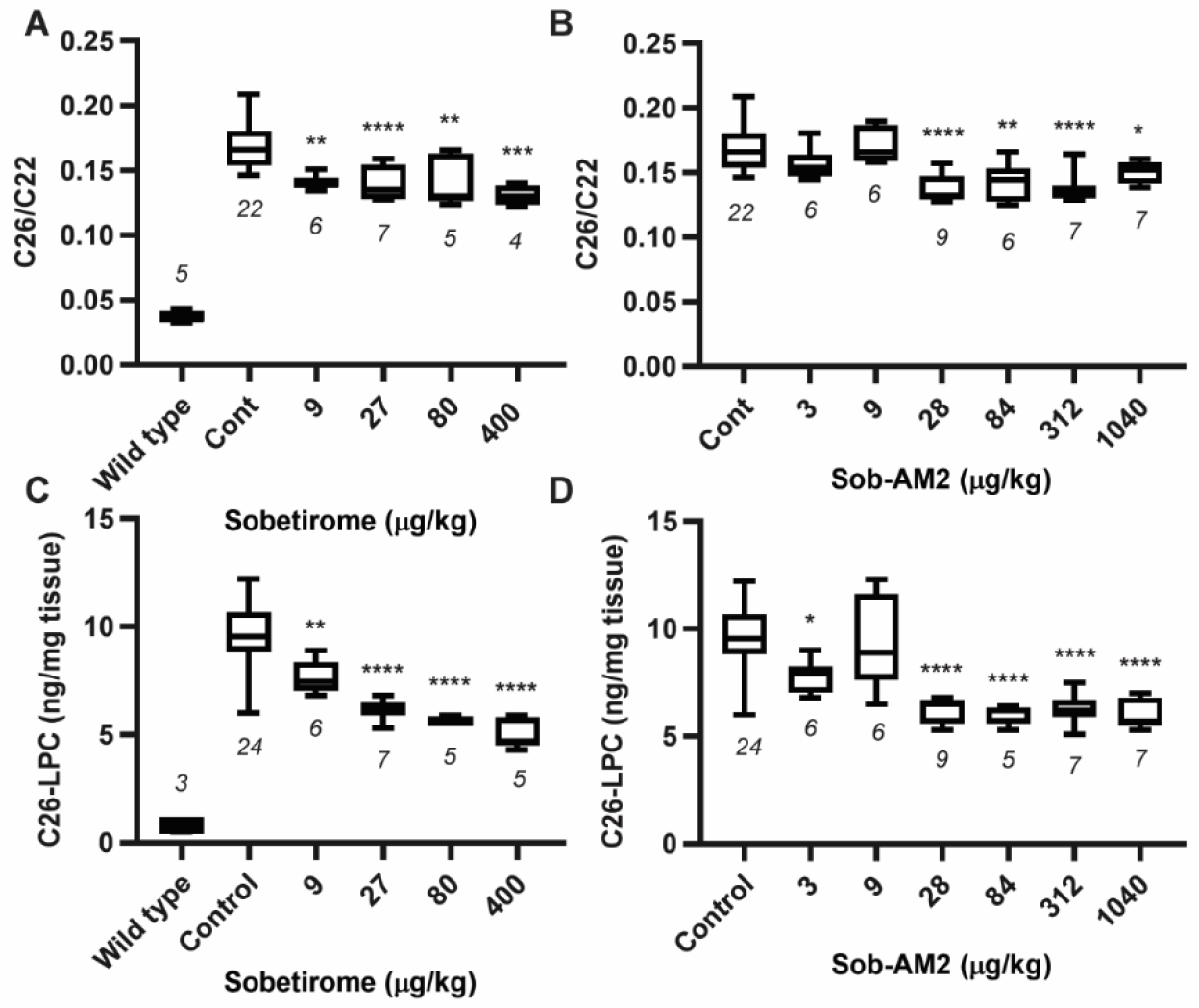
Sobetirome and Sob-AM2 lower C26-LPC and C26/C22 in the spinal cord. (A – D) Male *Abcd1* KO mice were administered chow containing sobetirome or Sob-AM2 from 3-15 weeks of age. The chow was compounded with sobetirome or Sob-AM2 at the estimated concentration required to administer the daily dose shown in the figure. (A and B) Total C26 and C22 in the spinal cord were measured by GC-MS and the C26/C22 ratio is reported. (C and D) C26-lysophosphatidylcholine (C26-LPC) in the spinal cord was measured by LC-MS/MS. All controls for each assay are combined into a single group. Wild-type mice levels (A and C) were not included in the statistical analyses. All data are represented as box and whisker plots with the error bars representing minimum and maximum. The number of animals is indicated below each dataset in the figure. Statistical analysis was performed using a one-way ANOVA test with Dunnett’s post-test for multiple comparisons between control and each dose (*P < 0.05, **P < 0.01, ***P < 0.001, ****P < 0.0001).

The oral bioavailability of Sob-AM2 is 20% in mice, while that of sobetirome is >90% (Ferrara et al., 2017; Meinig et al., 2019). To confirm that chow administration of either drug provides the anticipated sobetirome levels *in vivo*, sobetirome concentrations in serum and brain were measured in mice administered chow for 12 weeks at equimolar doses of 80 μg/kg sobetirome or 84 μg/kg Sob-AM2 (Devereaux et al., 2018; Placzek et al., 2016; Placzek and Scanlan, 2015). Serum levels of sobetirome were lower after Sob-AM2 administration (Figure 2C), and brain levels of sobetirome were similar from the two different drugs (Figure 2D). This arises from a combination of reduced Sob-AM2 oral bioavailability balanced by increased brain sobetirome exposure delivered from Sob-AM2. This is consistent with the C26/C22 and C26-LPC lowering observed in the CNS in which Sob-AM2 administration resulted in similar VLCFA reductions as sobetirome in the CNS (Figure 2B). Even though only 20% of Sob-AM2 was absorbed into systemic circulation as compared to 90% of sobetirome, the Sob-AM2 prodrug produces the same CNS effect, because it delivers more sobetirome to the CNS relative to peripheral exposure. Likewise, the reduced VLCFA lowering in serum and adrenal glands with an equal dose of Sob-AM2 versus sobetirome (Figure 2A) results from the reduced blood levels of sobetirome in mice that received Sob-AM2.

### Normalization of T4 levels increases lowering of VLCFAs in the periphery, but has no effect in the central nervous system

Chronic treatment with sobetirome or Sob-AM2 can lead to depletion of thyroid hormone levels by a mechanism involving central suppression of the HPT-axis (Ferrara et al., 2018). We confirmed that both sobetirome and Sob-AM2 deplete circulating T4 levels in *Abcd1* KO mice in a dose-dependent fashion (Figure 5A and 5B). This raised the question of whether systemically depleted thyroid hormone mitigated the VLCFA lowering effects of sobetirome. As evidence for this possibility, we observed an unusual bell-shaped dose-response curve for sobetirome with greater reductions in C26-LPC levels at 9 and 80 μg/kg than the intermediate dose 27 μg/kg in *Abcd1* KO mice (Figure 1A). All of these sobetirome doses and treatment durations were sufficient to significantly deplete systemic thyroid hormone levels (Figure 5A). We hypothesized that the attenuated VLCFA lowering observed in serum with 27 μg/kg sobetirome resulted from thyroid hormone depletion at this low sobetirome dose. This resulted in a lower total concentration of natural and unnatural thyroid hormone agonists, than 80 μg/kg or the 9 μg/kg dose, which only partially depleted endogenous thyroid hormone.

**Figure 5.**
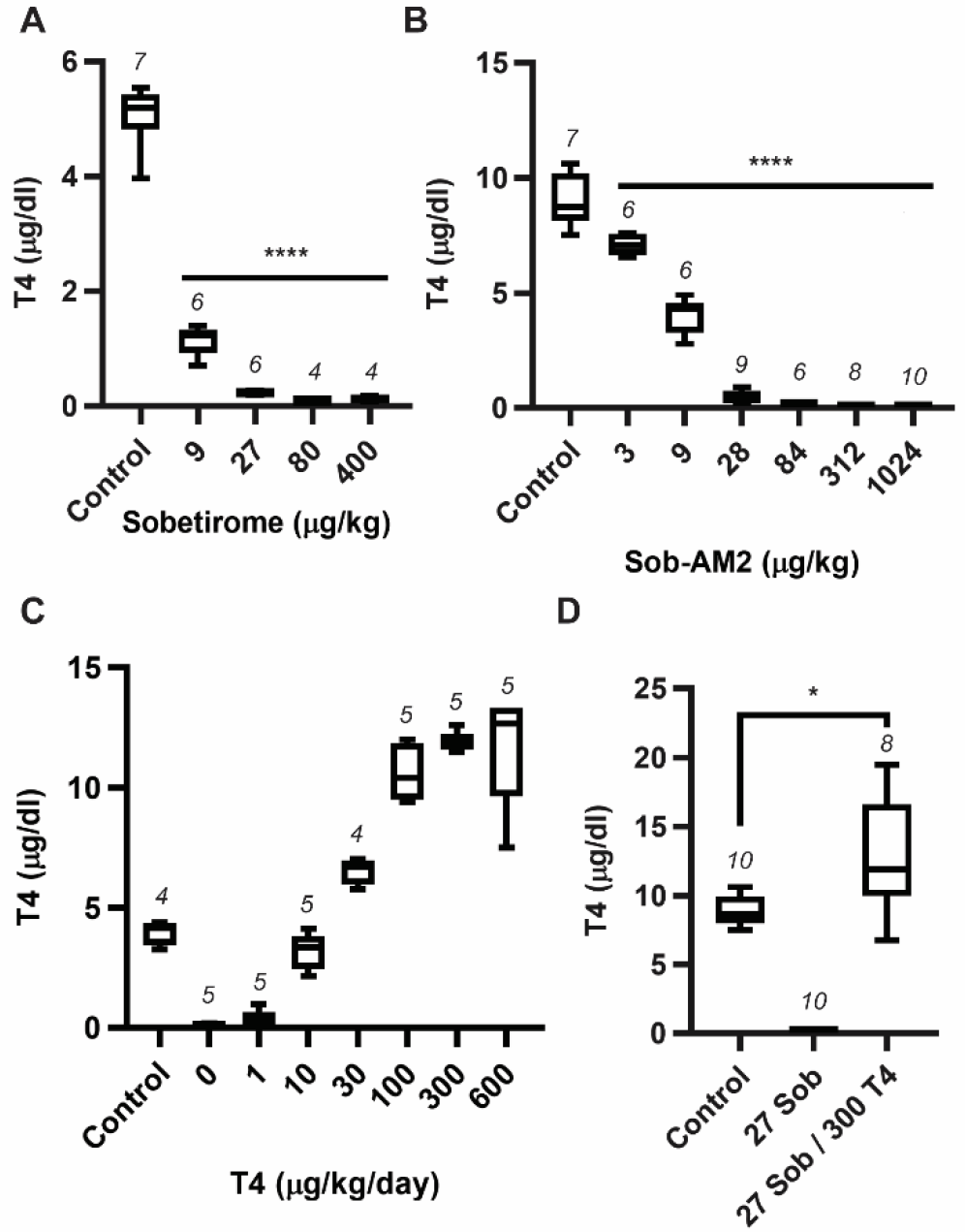
Sobetirome and Sob-AM2 suppress endogenous T4 levels, which can be restored by administration of T4. (A and B) Male *Abcd1* KO mice were administered chow containing sobetirome or Sob-AM2 from 3-15 weeks of age. The chow was compounded with sobetirome or Sob-AM2 at the estimated concentration required to administer the daily dose shown in the figure. Total T4 in serum was measured by radioimmunoassay. (C) Male *Abcd1* KO mice were administered chow containing sobetirome (27 μg/kg/day) for four weeks. After 4 weeks, the sobetirome chow was continued and the mice were co-administered daily T4 doses for 10 days by oral gavage. Total T4 in serum was measured by radioimmunoassay. (D) Male *Abcd1* KO mice were administered chow containing sobetirome (27 μg/kg/day) or chow containing sobetirome (27 μg/kg/day) and T4 (300 μg/kg/day) from 3-15 weeks of age. Total T4 in serum was measured by radioimmunoassay. All data are represented as box and whisker plots, and the error bars represent minimum and maximum. The number of animals is indicated above each dataset in the figure. Statistical analysis was performed using a one-way ANOVA test with Dunnett’s post-test for multiple comparisons between control and each dose (*P < 0.05, **P < 0.01, ***P < 0.001, ****P < 0.0001) for A and B and unpaired t-test for D (*P < 0.05).

To test the possibility that restoring thyroid hormone levels could further reduce VLCFAs in both the periphery and CNS, we determined the dose of oral T4 that, in combination with sobetirome administered in chow at 27 μg/kg, was required to restore T4 levels to normal (Figure 5C). Based on this dose response curve, we formulated a combination chow containing 27 μg/kg sobetirome and 300 μg/kg T4, and treated *Abcd1* KO mice for 12 weeks with the chow. The combination chow was well-tolerated by the mice and increased T4 levels slightly above the euthyroid reference range for this strain (Figure 5D). We observed that increased T4 levels resulted in more lowering of C26-LPC and C26/C22 in all peripheral tissues (Figure 6A-C). Restoring T4 increased VLCFA lowering in serum at the 27 μg/kg dose, which was consistent with our hypothesis. However, restoring T4 levels did not enhance lowering of either C26-LPC or C26/C22 in the CNS (Figure 6D-G).

**Figure 6.**
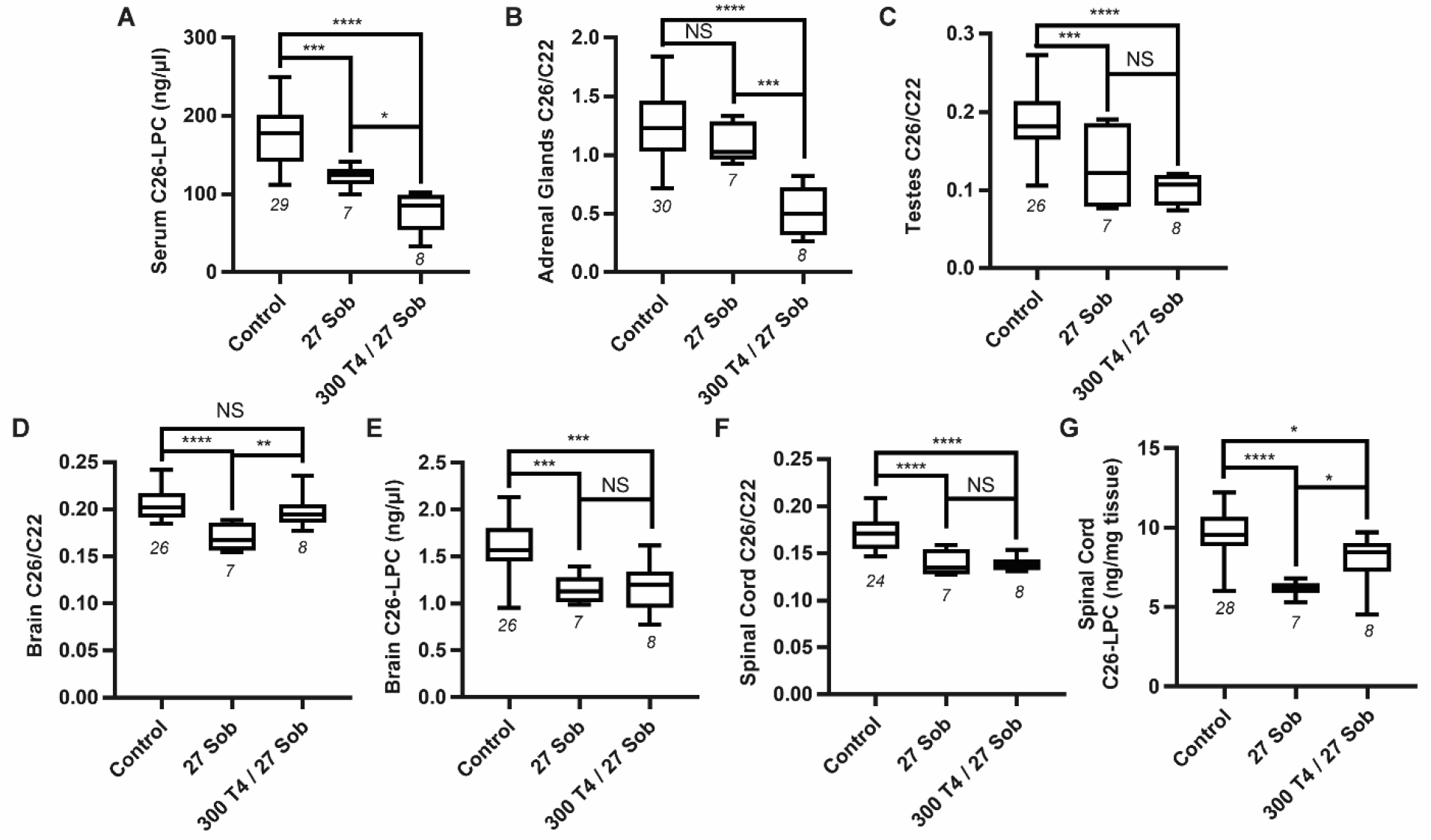
Co-administration of T4 and sobetirome enhances C26 lowering in peripheral tissues, but not in the CNS. Male *Abcd1* KO mice were administered chow containing sobetirome (27 μg/kg/day) or chow containing sobetirome (27 μg/kg/day) and T4 (300 μg/kg/day) from 3-15 weeks of age. C26-lysophosphatidylcholine (C26-LPC) was measured by LC-MS/MS in serum (A), brain (E), and spinal cord (G). Total C26 and C22 were measured by GC-MS and the C26/C22 ratio is reported for adrenal glands (B), testes (C), brain (D), and spinal cord (F). The number of animals is indicated below each dataset in the figure. All data are represented as box and whisker plots, and the error bars represent minimum and maximum. Statistical analysis was performed using a one-way ANOVA test with Tukey’s post-test for multiple comparisons between all groups (*P < 0.05, **P < 0.01, ***P < 0.001, ****P < 0.0001). NS indicates comparison found to be not significant.

### Aged *Abcd1* KO mice appear disease free without clinical signs similar to X-ALD

Male *Abcd1* KO mice have loss of function in Abcd1 similar to X-ALD patients and, like the patients, have elevated VLCFAs in circulation and all tissues (Forss-Petter et al., 1997; Kobayashi et al., 1997; Lu et al., 1997). However, at no point in their lifespan do *Abcd1* KO mice develop demyelinating brain lesions or disability resembling that which occurs in cerebral X-ALD patients. There is a report that describes a motor disability in *Abcd1* KO mice that are near the end of their natural lifespan (20 months) reminiscent of the disability that occurs in adrenomyeloneuropathy, the most common clinical phenotype of X-ALD. This motor disability in aged *Abcd1* KO mice was detected by a reduction in performance in two motor tests that correlated with histopathological abnormalities in spinal cord myelin (Pujol et al., 2002). In an effort to replicate this model, we performed monthly rotarod testing on *Abcd1* KO mice and wild-type littermates starting at 15 months of age. In the original study, a pronounced decline in rotarod latency was observed at 20 months in *Abcd1* KO compared to wild-type mice; however, we did not observe any rotarod performance decline or other evidence of motor disability in *Abcd1* KO or wild-type littermates at 20 months (Figure 7A). We followed a smaller cohort of *Abcd1* KO mice to 24 months, but still did not observe any decline in rotarod performance (Figure 7A). We also performed open field testing at 15 and 20 months, which in the original study revealed significant reductions in mobility and rearing events at both 15 and 20 months. However, we did not observe any differences in these parameters between wild-type and *Abcd1* KO mice at either timepoint (Figure 7B and 7C).

**Figure 7.**
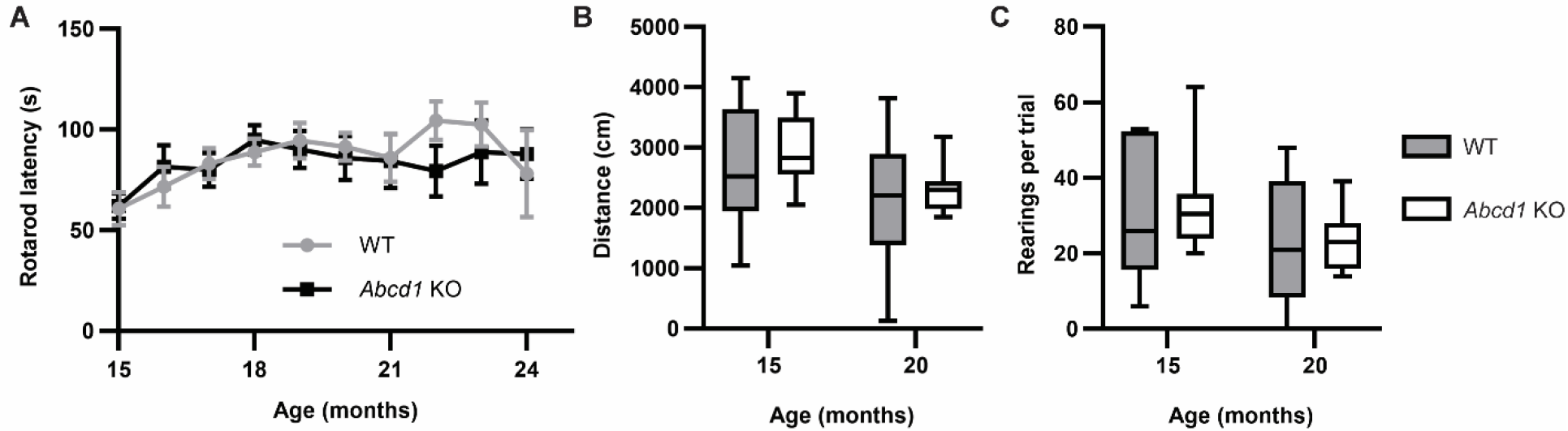
Aged *Abcd1* KO mice do not show motor deficits. Male *Abcd1* KO mice were aged to 20-24 months. (A) Starting at 15 months, the mice underwent monthly rotarod testing using the following program: 0-4 minutes with steady ramping from 8-40 RPM, and 4-5 min with the speed held at 40 RPM. Each test consisted of 3 trials with at least 15 minutes in between each trial, and the average latency was recorded.

From months 15-20, n = 10 for both wild-type (WT) and *Abcd1* KO; from months 21-24, n = 5 for both WT and *Abcd1* KO. Data are represented as mean latency, and the error bars represent SEM. (B and C) Open field testing was performed by videotaping each mouse for 10 minutes in a 40 cm x 40 cm white box. (B) Total distance traveled by each mouse was determined by tracking with EthoVision. (C) Total number of rearing events was determined by manual counting each video. For B and C, the data represent n = 10 for both WT and *Abcd1* KO. Statistical analysis performed using an unpaired t-test to compare WT and *Abcd1* KO showed no statistical difference at all timepoints in rotarod and open field testing.

## Discussion

The amount of VLCFA lowering in the periphery and CNS that is sufficient for therapeutic benefit in X-ALD is currently unknown, and would provide critical insight for development of X-ALD therapeutics. A robust translational animal model linking elevated VLCFAs to disease pathology similar to that occurs in X-ALD patients would be useful in this regard. A motor disability has been described for aged *Abcd1* KO mice reported as a fairly pronounced ~10-fold decrease in both rotarod latency and the frequency of rearing events in 20-month old mice, together resembling the myelopathy seen in adult AMN patients (Pujol et al., 2002) We were unable to reproduce these findings in our aged *Abcd1* KO mice, which were derived from a different engineered mouse line (Forss-Petter et al., 1997). Another report describes difficulty reproducing the rotarod disability in aged *Abcd1* KO mice, but instead discovers a more mild impairment to hind limb reflex extension in the aged mutant mice (Dumser et al., 2007). Other studies using aged *Abcd1/Abcd2* double KO mice have described a mild locomoter disability that can be partially corrected with PPAR agonist treatment (Morato et al., 2013); however, the double KO mice are not useful for evaluating VLCFA lowering therapies such as thyromimetics that stimulate *Abcd2* expression as a mechanism of action. In our hands *Abcd1* KO mice remain a reliable model for assessing efficacy of VLCFA lowering therapies in tissues relevant to X-ALD, but have limited utility for establishing that the observed VLCFA lowering is likely to be disease modifying for X-ALD.

Clinical data may provide insight into the amount of VLCFA lowering that is needed for a disease modifying therapy. Treatment with Lorenzo’s oil lowered VLCFAs in blood of X-ALD patients by ~50%, and it was suggested that it may delay onset of disease symptoms in asymptomatic children (Moser et al., 2007; Moser et al., 2005). However, Lorenzo’s oil does not appear to alter the progression of existing neurological symptoms in cerebral X-ALD patients, and it has been suggested that this is due to insufficient CNS exposure of the agent from a systemic dose (Berger et al., 2010). It is known that female carriers with one defective and one normal copy of *ABCD1* have elevated circulating VLCFA levels that are ~50% as elevated as male X-ALD patients. These heterozygous female carriers have mild neurological symptoms that develop much later in life than those of male patients, and do not typically present with the fatal brain involvement characteristic of cerebral ALD (Moser et al., 1983). Thus, a reasonable starting point for a disease modifying X-ALD therapy based on VLCFA lowering may be one that reduces the elevated circulating and CNS VLCFA levels by ~50%, which may be sufficient to prevent the development of childhood cerebral ALD, and delay the onset and reduce the severity of AMN.

We previously identified the maximum tolerated dose in mice of sobetirome-compounded chow to be 80 μg/kg/day for a 90-day dosing duration, which lowered VLCFAs in *Abcd1* KO mice by 15-20% (Hartley et al., 2017). Higher doses (including 400 μg/kg/day) caused weight loss starting around week-8 of a 12-week drug course, and this weight loss was usually a precursor to more serious adverse effects. Thus, we were previously unable to safely assess whether increased thyroid hormone action using sobetirome would further lower CNS VLCFAs. With the development of Sob-AM2, a prodrug of sobetirome with increased CNS penetration, we were able to increase the dose to 1040 μg/kg/day Sob-AM2 without any observed weight loss over a 12-week dosing period. Sob-AM2 has lower oral bioavailability than sobetirome, and a significant fraction of an oral Sob-AM2 dose is converted to sobetirome in mice by first-pass GI hydrolase activity (Meinig et al., 2019). However, even with these considerations, higher systemic exposure of Sob-AM2 was better tolerated by mice compared to an equivalent system exposure of sobetirome, presumably because the prodrug Sob-AM2 delivers less of the parent drug sobetirome to the blood (Figure S3A) from the combination of hydrolase activity in the CNS and periphery(Meinig et al., 2017). In the CNS, we observed that 12 weeks of Sob-AM2 treatment induced up to 40% lowering of C26-LPC and 15-20% lowering of total C26:0 with no weight loss. We have also shown recently in a separate study that sobetirome and Sob-AM2 are effective myelin repair agents (Hartley et al., 2019). This occurs through a mechanism involving thyromimetic induced oligodendrogenesis which is a potentially complementary therapeutic effect to VLCFA lowering in X-ALD. Sob-AM2 therefore has a superior thyromimetic profile that may be beneficial for treating all clinical phenotypes of X-ALD by correcting the biochemical abnormality in lipid metabolism and stimulating myelin repair. Furthermore, the potential of Sob-AM2 as an X-ALD development candidate is likely better than that of sobetirome due to the greater CNS penetration of Sob-AM2 with lower peripheral parent drug exposure and potential toxicity.

Sobetirome and Sob-AM2 induced dose-dependent central suppression of the HPT-axis in *Abcd1* KO mice, resulting in systemic depletion of thyroid hormone. We tested whether coadministration of sobetirome and T4 further enhanced CNS VLCFA lowering by replacing the depleted thyroid hormone, thus increasing the concentration of total—i.e. endogenous and synthetic—thyroid hormone agonists. Peripheral VLCFAs in serum, adrenal glands, and testes were further lowered by thyroid hormone replacement. However, sobetirome-induced VLCFA lowering in the CNS was unaffected by thyroid hormone replacement, and VLCFA lowering was similar to that observed with the sobetirome alone. These data suggest that the kinetics of VLCFA turnover, which regulates the timing and frequency of VLCFA transport to the peroxisome for degradation, is slower in the CNS than the periphery. This likely plays a role in the observable limits to CNS VLCFA lowering in *Abcd1* KO mice.

A question that arises is whether the slow kinetics of CNS VLCFA turnover would be altered in a demyelinating disease state similar to X-ALD. Since much of the CNS lipid inventory resides in myelin, it is conceivable that a demyelinating disease would liberate fatty acid derivatives such as VLCFAs from the lipid-rich environment of myelin. This disruption may increase VLCFA transport rate to peroxisomes, thereby increasing turnover and decreasing VLCFA half-life *in vivo.* For the development of therapeutics targeting mechanisms of lipid metabolism, it is critical to understand whether and how lipid turnover changes in a demyelinating disease state.

Thyromimetics induced greater lowering for C26-LPC as compared to the total C26:0 lipid population, which raises a second question concerning the pathogenicity of specific VLCFA derivatives such as C26-LPC in X-ALD. C26-LPC has emerged as the most important VLCFA in diagnosing X-ALD largely due to its compatibility with the LC-MS/MS bioanalytical techniques commonly used to identify and quantify lipids from biological matrices. Whether or not C26-LPC is more important than other C26 fatty acid derivatives in initiation or progression of X-ALD pathology remains an open question. Studies have shown that in brain tissue from X-ALD patients, C26:0 is present in all of the major classes of lipids, including glycerophospholipids, sphingolipids, and neutral lipids including cholesteryl esters (Brown et al., 1983; Wilson and Sargent, 1993). If a subset of the total C26:0 lipid population or a specific derivative such as C26-LPC could be identified as the major pathogenic variant, then specific targeting of this variant could be envisioned, which would accelerate the development of a therapeutic intervention in X-ALD.

### Significance

Elevated VLCFAs are linked to disease pathology in X-ALD, and therapeutic strategies for lowering VLCFAs may be beneficial for patients with X-ALD. This study demonstrates that the thyromimetics sobetirome and Sob-AM2 lower VLCFAs in the periphery by up to 60% in *Abcd1* KO mice. In the CNS, C26/C22 was lowered by 15-20% and C26-LPC by 25-40%, which appear to represent thresholds beyond which increased thyroid hormone agonism does not increase VLCFA lowering. This likely results from the slow kinetics of lipid turnover in *Abcd1* KO mice, which are more tolerant of systemically elevated VLCFA and do not display the characteristic clinical signs or pathology typical of symptomatic X-ALD male patients. The CNS-penetrating prodrug Sob-AM2 was more potent than sobetirome in lowering CNS VLCFAs in *Abcd1* KO mice, and was better tolerated due to lower peripheral parent drug exposure. The results of this study support the clinical development of CNS-penetrating thyromimetic prodrugs for VLCFA lowering, which is a promising therapeutic strategy for treating all clinical phenotypes of X-ALD.

## Acknowledgements

The research was supported by the National Institutes of Health (DK52798 to T.S.S.) and the Oregon Health & Sciences University (OHSU) Laura Fund for Innovation in Multiple Sclerosis (T.S.S.). M.D.H. received postdoctoral funding from the National Institutes of Health (2T32DK007680-21) and the National Multiple Sclerosis Society (FG 2023A 1/2) with partial support from the Dave Tomlinson Research Fund. Assistance with animal behavior studies was provided by Laura Villasana, Mike Jacobson and Helen Liu in the Department of Anesthesiology & Perioperative Medicine at OHSU. Analytical support was provided by the Bioanalytical Shared Resource/Pharmacokinetics Core Facility, which is part of the University Shared Resource Program at OHSU. We thank Lisa Bleyle for her assistance with both GC/MS and LC-MS/MS analysis. We would also like to thank Arjun Subramanian and Andrés Olavarrieta for their technical assistance.

## Author Contributions

Conceptualization: M.D.H., M.D.S., and T.S.S.; Methodology: M.D.H., M.D.S., M.J.D., and T.S.S.; Formal Analysis: M.D.H., M.D.S, and M.J.D.; Investigation: M.D.H., M.D.S, M.J.D., T.B., and L.L.K.; Writing – Original Draft: M.D.H. and T.S.S.; Writing – Review & Editing: M.D.H., M.D.S., M.J.D., and T.S.S.; Visualization: M.D.H.; Supervision: M.D.H. and T.S.S.; Funding Acquisition: M.D.H. and T.S.S.

## Declaration of Interests

T.S.S. is a founder of Llama Therapeutics, Inc. M.D.H. is a consultant to Llama Therapeutics, Inc. T.S.S. and M.D.H. are inventors on licensed pending and issued patents.

## STAR Methods

### Lead contact and materials availability

Further information and requests for resources and reagents should be directed to and will be fulfilled by the Lead Contact, Thomas S. Scanlan.

### Experimental models and subject details

#### Animal models

All experiments were performed with the approval of the Oregon Health & Science University (OHSU) IACUC committee. The *Abcd1* KO mice were reconstituted from embryos by Jackson Laboratory (strain #003716) in a C57BL6/J background. All mice in the described experiments were bred at OHSU and were housed under standard 12h/12h light/dark conditions. Mice were group housed with littermates at all times with no more than five mice per cage. Only male *Abcd1* KO mice were used in the experiments, as *Abcd1* is an X-linked gene, and X-ALD is a disease that primarily affects males. At weaning, tail tips were collected from the mice, and PCR was used to determine the genotype of the mice following protocols recommended by Jackson Laboratory.

### Method details

#### Animal experiments

For the 12-week treatment experiments, male pups were weaned at 3 weeks, and each cage containing wild-type and *Abcd1* KO littermates was randomly assigned to control or drug chow treatment groups until the experiments were fully enrolled (n = 4-8 per group). From 3-15 weeks of age, the mice were fed *ad lib* control chow (Envigo Teklad 2016) or control chow compounded with sobetirome or Sob-AM2 at the described doses. Sobetirome chow was prepared containing 0.04 and 0.13 mg/kg chow, and Sob-AM2 chow was prepared containing 0.02, 0.05, 0.14, 0.42, 1.56, and 5.12 mg/kg chow. All mice in the same cage received the same chow. To minimize the effects of variability between litters, all dosing groups contained mice from at least two different litters. If a single litter had a large number of male mice (>4), those mice were separated into multiple cages and placed on different chow treatments.

At 15 weeks of age, mice were euthanized with carbon dioxide followed by cervical dislocation. Blood, brain, spinal cord, testes, and adrenal glands were promptly harvested. Brain, spinal cord, and testes were frozen immediately and stored at −80 °C. Blood was collected in a microcentrifuge tube and allowed to clot on ice for at least 20 minutes. Clotted blood was spun in a microcentrifuge for 15 minutes at 7500 RPM, and serum was collected and stored at −80 °C. Adrenal glands were removed and stored in RNALater for up to one month prior to microdissection of the adrenal gland from the surrounding fat. Following microdissection, adrenal glands were stored at −80 °C.

To determine the T4 replacement dose, mice were administered sobetirome chow (0.13 mg/kg chow) for four weeks to induce central hypothyroidism. After four weeks, the mice were continued on the sobetirome chow and additionally received (or were administered) T4 via oral gavage daily for 10 days at a range of doses (1 – 600 μg/kg). The mice were euthanized 8 h after the final administration of T4, and serum and brain were collected as described above. Based on the data in these experiments, a combination chow was prepared with sobetirome (0.13 mg/kg chow) and T4 (1.5 mg/kg chow). The mice were administered the combination chow for 12 weeks and tissue collection was performed as described above.

For the motor behavior experiments, male mice were group housed and aged until 15 months. At 15 months, rotarod testing was initiated and performed monthly until 20 or 24 months of age. Rotarod testing was performed using the Roto-rod (IITC Life Science), in which latencies were automatically recorded by mice hitting sensors on the floor. A test comprised 3 five-minute trials, which were performed with at least 15 minutes in between trials, and the average was used as the latency. The program used was the following: a steady ramp of 8-40 RPM from 0:00 to 4:00, and a constant speed of 40 RPM held from 4:00 to 5:00. If mice rotated once around holding onto the bar, that was counted as a fall and the time was recorded. If the mice made it 5:00, then the latency was recorded as 300 sec, and they were removed from the rotarod. Several days (2-4) prior to the initial round of rotarod testing (at 15 months age), the mice were trained on the rotarod with 3 trials of 2 minutes each at 8 RPM. If mice fell off during training, they were placed back on the rod.

Open field testing was performed at 15 and 20 months of age. Each test was performed by placing mice in an open field box (40 cm x 40 cm) and video recording their movements during a 10-minute period. The animal’s movements were analyzed using EthoVision to determine the total distance travelled during the 10-minute test. In addition, the total number of rearing events was determined manually by watching each video. Rearings were defined as events where mice were clearly standing on hind legs either with or without support from the wall.

#### T4 assay

T4 levels were determined by radioimmunoassay following the manufacturer protocol (IVD Technologies). Standards (0 – 10 μg/dl) or sample serum (25 μl) and tracer ^125^I-T4 (200 μl) were added to T4 antibody-coated tubes. The tubes were vortexed and incubated for two hours at room temperature. The liquid was decanted, and the tubes were washed twice with 1 ml of deionized water. The radioactivity (CPM) was determined with a gamma counter, and a standard curve was prepared by plotting log (concentration of ^125^I-T4 standards) vs. % bound (sample CPM/blank CPM x 100).

#### LC-MS/MS quantification of C26-LPC

Serum samples (10 μl) were vortexed after the addition of 1 ng of d_4_C26-LPC internal standard in methanol. The samples were extracted in methanol (130 μl) with vortexing. After incubation (10 min, room temperature (RT)), debris was pelleted, and the supernatant was analyzed via LC-MS/MS (Hubbard et al., 2009; Hubbard et al., 2006; Sandlers et al., 2012). Whole brain was homogenized at 200 mg tissue/ml in water, and spinal cord was homogenized at 100 mg/ml in water using a BeadMill 24 (ThermoScientific) with 3 metal beads per 2 ml tube. Internal standard (20 ng of d_4_C26-LPC) was added to 1 ml of diluted homogenate (4 mg/ml for brain, 1 mg/ml for spinal cord) and thoroughly vortexed. The samples were extracted in 2 ml of 1:1 butanol:0.5 M hydrochloric acid with vortexing and incubation (15 min, RT). Samples were separated by centrifugation (3000 x g, 10 min), and the butanol layer was removed and dried under vacuum. The final dried sample was dissolved in 150 μl of 10% dimethylformamide in methanol and analyzed by LC-MS/MS.

An Applied Biosystems 5500 QTRAP hybrid/triple quadrupole linear ion trap mass spectrometer was utilized to detect C26:0-LPC samples in positive mode with electrospray ionization using multiple reaction monitoring (MRM). Chromatographic separation was achieved over an 8 min analysis period using a Thermo Scientific BetaBasic C8 or C18 column (20 x 2.1 mm, 5 μm particle size). The solvent system comprised mobile phase A (0.028% ammonium hydroxide in water) and mobile phase B (0.028% ammonium hydroxide in isopropanol). The working gradient began with 40% mobile phase B, which was increased to 98% mobile phase B over 5 min, held until 6 min, and then returned to 30% organic by 6.1 min and held until the end of the analysis. The injection volume was 10 μl, and the solvent flow rate was 0.5 ml/min. The column temperature was held at 40 °C throughout the experiment. The instrument parameters for the MRM transitions in positive mode were optimized by direct infusion for the following transitions: C26-LPC, *m/z* 636 → 184 and *m/z* 636 → 104; d_4_C26-LPC, *m/z* 640 → 184 and *m/z* 640 → 104. Peaks were analyzed with Analyst 1.6.2 (Sciex) software. A linear standard curve was prepared by spiking standard and internal standard into the appropriate matrix from wild-type mice. For serum, the standard curve ranged from 1.25-1250 pg/μl. For brain and spinal cord, the standard curve ranged from 0.25-50 ng/mg tissue. The data in Figures 1–4 and 6 were normalized by tissue weight in the assay.

#### GC-MS quantification of total VLCFAs

Sample preparation was performed following published protocols (Lagerstedt et al., 2001). Tissues were homogenized in water using a BeadMill 24 (ThermoScientific). Brain was homogenized at 200 mg/ml, spinal cord at 100 mg/ml, and testes at 100 mg/ml. Prior to the assay, spinal cord homogenate was diluted to 10 mg/ml. Adrenal glands (2 per mice) were homogenized in 1 ml of water per adrenal gland. Tissue homogenates (25 μl of brain, 100 μl of diluted spinal cords, 400 μl of adrenal glands, or 200 μl of testes) were extracted in the presence of internal standards (d_4_C22:0, d_4_C24:0, and d_4_C26:0) in 2 ml of 2:3 isopropanol:hexane for 1 hr at room temperature (RT) with shaking. For brain, 1.00 μg of d_4_C22:0 and 0.16 μg of d_4_C26:0 were added to each sample. For spinal cord, 1.50 μg of d_4_C22:0 and 0.2 μg of d_4_C26:0 were added to each sample. For testes, 0.25 μg of d_4_C22:0 and 0.05 μg of d_4_C26:0 were added to each sample. For adrenal glands, 0.05 μg of d_4_C22:0 and 0.05 μg of d_4_C26:0 were added to each sample. The extracted samples underwent acid hydrolysis (2 ml of 9:1 acetonitrile: 6 M hydrochloric acid) followed by base hydrolysis (additional 2 ml of 9:1 methanol:10 M sodium hydroxide). Both hydrolysis steps were performed in capped 16 x 100 mm tubes for 45 minutes at 100 °C. After re-acidification (350 μl of 6 M hydrochloric acid), samples were extracted with hexanes (2 ml), and the hexane layer was dried under vacuum. Dried samples underwent derivatization with PFB (pentafluorobenzyl bromide, 50 μl 9:1 dry acetonitrile:PFB) in the presence of triethylamine (10 μl) for 1 hour at RT and were extracted again in hexanes (1 ml with 150 μl of 0.1 M hydrochloric acid), and the hexane layer was dried under vacuum. The final sample was dissolved in 50 μl of hexanes and analyzed by GC-MS.

Sample analysis was performed using an Agilent-7890B/5977A GC/MS operating in negative chemical ionization mode with methane as the reagent gas. Peaks were obtained over a 20-minute analysis period for the PFB esterified fatty acids using an Agilent HP-5ms column (30 m x 0.25 mm; film 0.25 μm), with helium as the carrier gas. The working temperatures of the source and transfer line were 250 and 325 °C, respectively. The split/splitless injector was held at 275 °C and was operated with a 1:25 split. The sample injection volume was 1 μl. The initial oven temperature was 150 °C, with a ramp rate of 15 °C/min, and a final temperature of 325 °C, held for 7 min. Acquisition was performed in the selected ion-monitoring (SIM) mode, with a dwell time of 25 ms. The ion *m/z* values for the endogenous standards were 339.3, 367.4, and 395.4 for C22:0, C24:0, and C26:0, respectively, while the deuterated internal standards had *m/z* values of 343.4, 371.4, and 399.4 for d_4_C22:0, d_4_C24:0, d_4_C26:0, respectively. Peaks were analyzed with Masshunter (Agilent) software. Standard calibration curves were generated based on the peak area ratios of each fatty acid matched to the deuterated internal standard. For brains, the standard curves ranged from 50-1250 ng/mg tissue for C22:0 and 4-100 ng/mg tissue for C26:0. For spinal cord, the standard curves ranged from 250-500 ng/mg tissue for C22:0 and 37.5-750 ng/mg tissue for C26:0. For adrenal glands, the standard curves ranged from 12.5-625 ng/adrenal gland for C22:0 and C26:0. For testes, the standard curves ranged from 0.75-37.5 ng/mg tissue for C22:0 and 0.13-6.25 ng/mg tissue for C26:0. All values were reported as C26/C22 ratios.

#### LC-MS/MS quantification of sobetirome

Serum samples (25 μl) were vortexed after the addition of 0.25 pmoles of d_6_-sobetirome internal standard (Placzek and Scanlan, 2015) in 10 μl 1:9 DMSO:water. The samples were extracted in acetonitrile (75 μl) with vortexing. Debris was removed with centrifugation at 10,000 x g for 15 minutes, and the supernatant was analyzed via LC-MS/MS (Devereaux et al., 2018; Placzek et al., 2016). Whole or half brain was homogenized at 200 mg tissue/ml in water using a BeadMill 24 (ThermoScientific) with 3 metal beads per 2 ml tube. Internal standard (0.15 pmoles of d_6_-sobetirome in 1:9 DMSO:water) was added to 125 μl of homogenate and thoroughly vortexed. The samples were extracted in 500 μl acetonitrile with vortexing. Debris was removed with centrifugation at 10,000 x g for 15 minutes, and the organic layer was removed and dried under vacuum. The final dried sample was dissolved in 60 μl of 1:1 acetonitrile:H2O and analyzed by LC-MS/MS.

An Applied Biosystems 5500 QTRAP hybrid/triple quadrupole linear ion trap mass spectrometer was utilized to detect C26:0-LPC samples in positive mode with electrospray ionization using multiple reaction monitoring (MRM). Chromatographic separation was achieved over an 8 min analysis period using a Hamilton PRP-C18 column (50 x 2.1 mm, 5 μm particle size). The solvent system comprised mobile phase A (10 mM ammonium formate in water) and mobile phase B (10 mM ammonium formate in 9:1 acetonitrile:water). The initial concentration of mobile phase B was 10% held for 0.5 min. The gradient of mobile phase B increased from 10% to 98% over 4.6 min, was held 1.9 min, and then returned to 10% organic by 7.1 min and held until the end of the analysis. The injection volume was 10 μl, and the solvent flow rate was 0.5 ml/min. The column temperature was held at 40 °C throughout the experiment. The instrument parameters for the MRM transitions in positive mode were optimized by direct infusion for the following transitions: sobetirome, *m/z* 327.3 → 269.3, 327.3 → 269.0, and *m/z* 327.3 → 135.0; dβ-sobetirome, *m/z* 333.0 → 275.2 and *m/z* 333.0 → 141.1. Peaks were analyzed with Multiquant software. A linear standard curve was prepared by spiking standard and internal standard into the appropriate matrix from wild-type mice. For serum, the standard curve ranged from 0.1-1000 ng/ml. For brain, the standard curve ranged from 0.1-100 ng/g tissue.

#### Quantification and statistical analysis

Statistics were performed by unpaired t-test or one-way ANOVA. The Dunnett’s or Tukey’s posttests for multiple comparisons were used to determine statistical significance as indicated in the figure legend. Significance is indicated in each figure with asterisks (*P < 0.05, **P < 0.01, ***P < 0.001, and ****P < 0.0001). All controls were combined into a single group. The ROUT outlier test was used to identify any statistical outliers, and those samples were excluded from the data. The following groups had a single outlier identified by the ROUT outlier test that was excluded from the data. Figure 1: serum Sob-AM2 (9 μg/kg), testes control, testes Sob-AM2 (1040 μg/kg), and adrenal glands control; Figure 3: brain C26/C22 control, brain C26-LPC SobAM2 (84 μg/kg), and brain C26-LPC Sob-AM2 (312 μg/kg); and Figure S3: brain sobetirome in SobAM2 (84 μg/kg). In all experiments, the n indicates the number of mice, and the exact n (excluding outliers) is provided in the figures, figure legends, and Tables S1 and S2.

#### Data and code availability

This study did not generate any datasets or code.

## Notes

#### Summary of Updates

Supplemental tables added to include the absolute values for all C26-LPC, C22:0, and C26:0 measurements; Fig. 2, 5, and S3 combined into new Fig. 2; Revised discussion of Fig. 7; Wild-type values added for each tissue.

