## Supplemental Figures and Tables for "Pharmacological complementation remedies an inborn error of lipid metabolism"

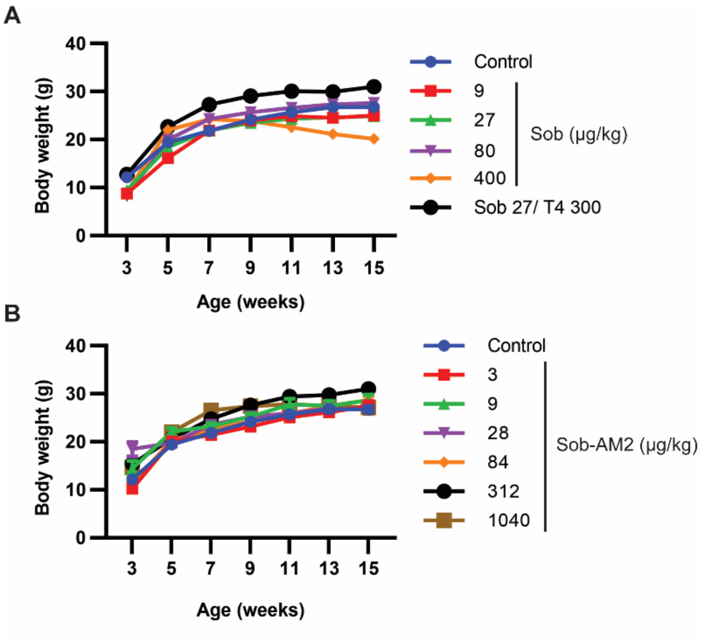


**Figure S1 (related to Figures 1-5). Body weights during sobetirome and Sob-AM2 dosing.**

Male *Abcd1* KO mice were administered chow containing sobetirome or Sob-AM2 from 3-15 weeks of age. The chow was compounded with (A) sobetirome or (B) Sob-AM2 at the concentration required to administer the estimated daily dose shown in the figure. Body weights were recorded every two weeks. All data are represented as the mean, and the error bars represent SEM. Only mice administered sobetirome at 400 µg/kg showed significant weight loss during the 12-week dosing period.

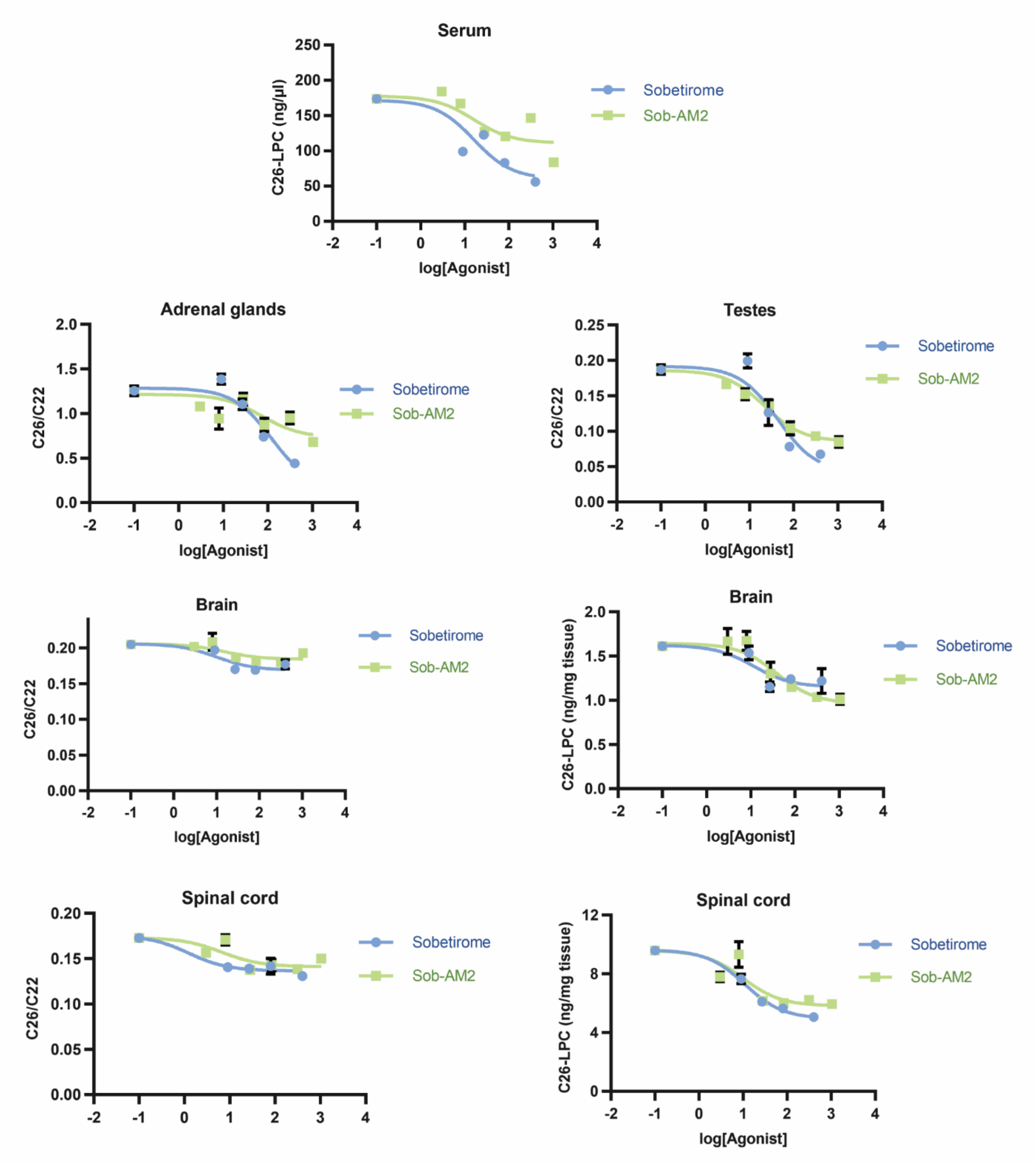


**Figure S2 (related to Figures 1-5). Dose-response curves for sobetirome and Sob-AM2.**

Male *Abcd1* KO mice were administered chow containing sobetirome or Sob-AM2 from 3-15 weeks of age. The chow was compounded with sobetirome or Sob-AM2 at the concentration required to administer the estimated daily dose shown in the figure. C26-lysophosphatidylcholine (C26-LPC) was measured by LC-MS/MS in serum, brain, and spinal cord. Total C26 and C22 were measured by GC-MS and the C26/C22 ratio is reported for adrenal glands, testes, brain, and spinal cord. All data are represented as the mean, and the error bars represent SEM.

**Table S1 (related to Figures 1, 3, 4, and 7). C26-LPC values and percent change relative to control are indicated as mean ± SEM for serum, brain, and spinal cord.**

| Drug (µg/kg) | Serum C26-LPC (ng/ml) | % change relative to control | n | Brain C26-LPC (ng/mg tissue) | % change relative to control | n |
|---|---|---|---|---|---|---|
| Wild-type | 5.8 ± 0.9 | - | 4 | 0.09 ± 0.02 | | 4 |
| Control (vehicle) | 173.7 ± 6.4 | - | 29 | 1.61 ± 0.05 | - | 28 |
| Sobetirome (9) | 99.2 ± 5.7 | -43 ± 3 | 6 | 1.54 ± 0.08 | -5 ± 5 | 6 |
| Sobetirome (27) | 122.8 ± 5.2 | -29 ± 3 | 7 | 1.16 ± 0.05 | -28 ± 3 | 7 |
| Sobetirome (80) | 83.1 ± 4.1 | -52 ± 2 | 5 | 1.24 ± 0.03 | -23 ± 2 | 5 |
| Sobetirome (400) | 56.2 ± 6.3 | -68 ± 4 | 5 | 1.22 ± 0.14 | -24 ± 9 | 4 |
| Sob-AM2 (3) | 184.3 ± 8.1 | 6 ± 5 | 6 | 1.61 ± 0.05 | 3 ± 9 | 6 |
| Sob-AM2 (9) | 167.4 ± 3.5 | -4 ± 2 | 5 | 1.67 ± 0.14 | 4 ± 7 | 6 |
| Sob-AM2 (28) | 127.1 ± 8.2 | -27 ± 5 | 9 | 1.67 ± 0.11 | -19 ± 8 | 9 |
| Sob-AM2 (84) | 120.4 ± 5.2 | -31 ± 3 | 6 | 1.30 ± 0.13 | -29 ± 2 | 6 |
| Sob-AM2 (312) | 146.9 ± 14.6 | -15 ± 8 | 7 | 1.15 ± 0.03 | -36 ± 3 | 7 |
| Sob-AM2 (1040) | 83.9 ± 5.0 | -52 ± 3 | 7 | 1.04 ± 0.05 | -37 ± 4 | 7 |
| Sobetirome (27) and T4 (300) | 78.1 ± 9.1 | -55 ± 5 | 8 | 1.19 ± 0.09 | -26 ± 6 | 8 |

| Drug (µg/kg) | Spinal cord C26-LPC (ng/mg tissue) | % change relative to control | n |
|---|---|---|---|
| Wild-type | 0.73 ± 0.07 | - | 3 |
| Control (vehicle) | 9.59 ± 0.29 | - | 24 |
| Sobetirome (9) | 7.65 ± 0.31 | -20 ± 3 | 6 |
| Sobetirome (27) | 6.11 ± 0.18 | -36 ± 2 | 7 |
| Sobetirome (80) | 5.64 ± 0.10 | -41 ± 1 | 5 |
| Sobetirome (400) | 5.06 ± 0.31 | -47 ± 1 | 5 |
| Sob-AM2 (3) | 7.80 ± 0.32 | -19 ± 3 | 6 |
| Sob-AM2 (9) | 9.33 ± 0.89 | -3 ± 9 | 6 |
| Sob-AM2 (28) | 6.17 ± 0.20 | -36 ± 2 | 9 |
| Sob-AM2 (84) | 6.02 ± 0.20 | -37 ± 2 | 5 |
| Sob-AM2 (312) | 6.23 ± 0.28 | -35 ± 3 | 7 |
| Sob-AM2 (1040) | 5.94 ± 0.25 | -38 ± 3 | 7 |
| Sobetirome (27) and T4 (300) | 8.00 ± 0.58 | -17 ± 6 | 8 |

**Table S2 (related to Figures 1 and 7). C22, C26, C26/C22, and percent change relative to control (mean ± SEM) for adrenal glands.**

| Drug (µg/kg) | n | C22 (ng/gland) | C26 (ng/gland) | C26/C22 | % change relative to control |
|---|---|---|---|---|---|
| Wild-type | 5 | 550 ± 69 | 130 ± 39 | 0.23 ± 0.07 | - |
| Control (vehicle) | 30 | 324 ± 27 | 376 ± 43 | 1.26 ± 0.05 | - |
| Sobetirome (9) | 6 | 551 ± 24 | 765 ± 52 | 1.38 ± 0.06 | 10 ± 5 |
| Sobetirome (27) | 7 | 574 ± 39 | 639 ± 63 | 1.11 ± 0.06 | -12 ± 5 |
| Sobetirome (80) | 5 | 226 ± 10 | 168 ± 12 | 0.74 ± 0.02 | -41 ± 2 |
| Sobetirome (400) | 5 | 288 ± 14 | 128 ± 13 | 0.44 ± 0.04 | -65 ± 3 |
| Sob-AM2 (3) | 6 | 549 ± 33 | 592 ± 42 | 1.08 ± 0.04 | -14 ± 4 |
| Sob-AM2 (9) | 5 | 346 ± 17 | 322 ± 35 | 0.94 ± 0.11 | -25 ± 9 |
| Sob-AM2 (28) | 9 | 827 ± 88 | 937 ± 84 | 1.16 ± 0.07 | -8 ± 5 |
| Sob-AM2 (84) | 6 | 1038 ± 155 | 856 ± 63 | 0.87 ± 0.07 | -31 ± 6 |
| Sob-AM2 (312) | 7 | 772 ± 83 | 718 ± 67 | 0.95 ± 0.07 | -24 ± 5 |
| Sob-AM2 (1040) | 7 | 713 ± 98 | 486 ± 70 | 0.68 ± 0.03 | -46 ± 2 |
| Sobetirome (27) and T4 (300) | 8 | 693 ± 97 | 360 ± 72 | 0.51 ± 0.08 | -59 ± 6 |

**Table S3 (related to Figures 1 and 7). C22, C26, C26/C22, and percent change relative to control (mean ± SEM) for testes.**

| Drug (µg/kg) | n | C22 (ng/mg tissue) | C26 (ng/mg tissue) | C26/C22 | % change relative to control |
|---|---|---|---|---|---|
| Wild-type | 5 | 1.30 ± 0.13 | 0.07 ± 0.01 | 0.062 ± 0.011 | - |
| Control (vehicle) | 26 | 1.22 ± 0.09 | 0.22 ± 0.02 | 0.187 ± 0.007 | - |
| Sobetirome (9) | 6 | 0.78 ± 0.01 | 0.16 ± 0.01 | 0.199 ± 0.010 | 7 ± 5 |
| Sobetirome (27) | 7 | 0.96 ± 0.06 | 0.12 ± 0.02 | 0.127 ± 0.018 | -32 ± 10 |
| Sobetirome (80) | 5 | 0.72 ± 0.04 | 0.06 ± 0.01 | 0.078 ± 0.004 | -58 ± 2 |
| Sobetirome (400) | 4 | 0.78 ± 0.09 | 0.05 ± 0.01 | 0.068 ± 0.003 | -64 ± 1 |
| Sob-AM2 (3) | 6 | 1.46 ± 0.05 | 0.24 ± 0.01 | 0.166 ± 0.005 | -11 ± 3 |
| Sob-AM2 (9) | 6 | 1.33 ± 0.06 | 0.20 ± 0.02 | 0.152 ± 0.008 | -19 ± 4 |
| Sob-AM2 (28) | 9 | 1.35 ± 0.06 | 0.18 ± 0.01 | 0.136 ± 0.008 | -27 ± 4 |
| Sob-AM2 (84) | 6 | 1.28 ± 0.06 | 0.13 ± 0.02 | 0.104 ± 0.009 | -44 ± 5 |
| Sob-AM2 (312) | 7 | 1.43 ± 0.07 | 0.13 ± 0.01 | 0.093 ± 0.006 | -50 ± 3 |
| Sob-AM2 (1040) | 6 | 1.54 ± 0.12 | 0.18 ± 0.05 | 0.085 ± 0.007 | -55 ± 4 |
| Sobetirome (27) and T4 (300) | 8 | 1.42 ± 0.06 | 0.14 ± 0.01 | 0.102 ± 0.007 | -45 ± 4 |

**Table S4 (related to Figures 3, 4, and 7). C22, C26, C26/C22, and percent change relative to control (mean ± SEM) for brains.**

| Drug (µg/kg) | n | C22 (ng/mg tissue) | C26 (ng/mg tissue) | C26/C22 | % change relative to control |
|---|---|---|---|---|---|
| Wild-type | 4 | 22.3 ± 2.4 | 0.8 ± 0.1 | 0.035 ± 0.004 | - |
| Control (vehicle) | 26 | 35.4 ± 1.2 | 7.2 ± 0.3 | 0.205 ± 0.003 | - |
| Sobetirome (9) | 6 | 38.1 ± 0.9 | 7.5 ± 0.2 | 0.197 ± 0.002 | -4 ± 1 |
| Sobetirome (27) | 7 | 35.8 ± 1.8 | 6.1 ± 0.3 | 0.170 ± 0.005 | -17 ± 3 |
| Sobetirome (80) | 5 | 39.7 ± 0.8 | 6.7 ± 0.2 | 0.169 ± 0.005 | -17 ± 2 |
| Sobetirome (400) | 4 | 40.3 ± 1.0 | 7.3 ± 0.3 | 0.182 ± 0.006 | -11 ± 3 |
| Sob-AM2 (3) | 6 | 38.5 ± 0.6 | 7.8 ± 0.2 | 0.202 ± 0.005 | -1 ± 2 |
| Sob-AM2 (9) | 6 | 35.0 ± 1.0 | 7.3 ± 0.5 | 0.209 ± 0.012 | 2 ± 6 |
| Sob-AM2 (28) | 9 | 40.8 ± 1.3 | 7.6 ± 0.2 | 0.187 ± 0.003 | -9 ± 1 |
| Sob-AM2 (84) | 5 | 37.0 ± 0.7 | 6.7 ± 0.1 | 0.182 ± 0.002 | -11 ± 1 |
| Sob-AM2 (312) | 5 | 28.6 ± 1.6 | 5.1 ± 0.3 | 0.180 ± 0.004 | -12 ± 2 |
| Sob-AM2 (1040) | 7 | 25.4 ± 1.6 | 4.9 ± 0.3 | 0.193 ± 0.005 | -6 ± 2 |
| Sobetirome (27) and T4 (300) | 8 | 26.1 ± 2.6 | 5.3 ± 0.7 | 0.199 ± 0.006 | -3 ± 3 |

**Table S5 (related to Figures 3, 4, and 7). C22, C26, C26/C22, and percent change relative to control (mean ± SEM) for spinal cords.**

| Drug (µg/kg) | n | C22 (ng/mg tissue) | C26 (ng/mg tissue) | C26/C22 | % change relative to control |
|---|---|---|---|---|---|
| Wild-type | 5 | 1184 ± 68 | 44 ± 3 | 0.037 ± 0.001 | - |
| Control (vehicle) | 22 | 1129 ± 23 | 191 ± 11 | 0.169 ± 0.003 | - |
| Sobetirome (9) | 6 | 1210 ± 23 | 170 ± 3 | 0.141 ± 0.002 | -17 ± 1 |
| Sobetirome (27) | 7 | 1198 ± 24 | 167 ± 7 | 0.139 ± 0.005 | -18 ± 3 |
| Sobetirome (80) | 5 | 1841 ± 161 | 257 ± 12 | 0.142 ± 0.009 | -16 ± 5 |
| Sobetirome (400) | 4 | 1361 ± 78 | 177 ± 5 | 0.131 ± 0.004 | -23 ± 2 |
| Sob-AM2 (3) | 6 | 1091 ± 64 | 170 ± 7 | 0.157 ± 0.005 | -7 ± 3 |
| Sob-AM2 (9) | 6 | 1059 ± 86 | 180 ± 14 | 0.171 ± 0.006 | 1 ± 3 |
| Sob-AM2 (28) | 9 | 1109 ± 28 | 152 ± 4 | 0.138 ± 0.004 | -19 ± 2 |
| Sob-AM2 (84) | 6 | 1173 ± 44 | 167 ± 6 | 0.143 ± 0.006 | -16 ± 4 |
| Sob-AM2 (312) | 7 | 1272 ± 44 | 176 ± 7 | 0.138 ± 0.005 | -18 ± 3 |
| Sob-AM2 (1040) | 7 | 1083 ± 36 | 162 ± 6 | 0.150 ± 0.003 | -11 ± 2 |
| Sobetirome (27) and T4 (300) | 8 | 1134 ± 68 | 158 ± 11 | 0.139 ± 0.003 | -18 ± 2 |
